# Genome-wide characterization and expression profiling of B3 superfamily during flower induction stage in pineapple (*Ananas comosus* L.)

**DOI:** 10.1101/2020.10.19.345181

**Authors:** Cheng Cheng Ruan, Zhe Chen, Fu Chu Hu, Xiang He Wang, Li Jun Guo, Hong Yan Fan, Zhi Wen Luo, Zhi Li Zhang

## Abstract

The B3 superfamily is a plant-specific family, which involves in growth and development process. By now, the identification of B3 superfamily in pineapple (*Ananas comosus*) has not reported. In this study, 57 *B3* genes were identified and further phylogenetically classified into five subfamilies, all genes contained B3 domain. Chromosomal localization analysis revealed that 54 of 57 *AcB3* genes were located on 21 chromosomes.Collinearity analysis indicated that the segmental duplication was the main event in the evolution of B3 gene superfamily and most of them were under purifying selection. Moreover, there were 7 and 39 pairs of orthologous B3s were identified between pineapple and *Arabidopsis* or rice, respectively, which indicated the closer genetic relationship between pineapple and rice. Most genes had *cis*-element of abscisic acid, ethylene, MeJA, light, and abiotic stress. qRT-PCR showed that the expression level of most *AcB3* genes were up-regulated within 1 d after ethephon treatment and expressed high level in flower bud differentiation period in stem apex. This study provide a comprehensive understanding of *AcB3s* and a basis for future molecular studies of ethephon induced pineapple flowering.

## Introduction

Transcription factor (TF) is a protein with special structure which regulating gene expression by binding to transcriptional regulatory regions, also known as trans-acting factors [1–2]. There are about 2000 TFs in *Arabidopsis thaliana*, they play the role mainly in biotic and abiotic stress response, organ development such as flowering, hormone signal response, cell growth and differentiation [3]. Flowering is an important process, which has a direct impact on the reproduction and yield of plants [4]. Many TFs are involved in flowering: *AP2*, *bHLH*, *bZIP*, *MADS* and *B3* [5–6].

The B3-type DNA binding domain (DBD) was first found in *VIVIPAROUS1* (*VP1*) from *Zea mays*, all members of the B3 superfamily contain this domain, and this kind of TF is only exist in plant [7–8]. The number of B3 superfamily gene differs in different plant, there are 118, 91, 81, 108 *B3* genes in *Arabidopsis thaliana*, *Oryza sativa*, *Zea mays*, *Glycine max*, respectively [9–10]. Base on the structure and function of protein, the B3 superfamily can be devided into five members: *auxin response factor* (*ARF*), *Leafy Cotyledon 2* (*LEC2*)-*Abscisic Acid Insensitive* 3(*ABI3*)-*VAL*(*LAV*), *high level expression of sugar inducible* (*HSI*), *related to ABI3/VP1* (*RAV*) and *reproductive meristem* (*REM*) [11–12]. In *Arabidopsis thaliana*, *ABI3* (one B3 domain), *HIS* (one B3 domain and zf-CW domain), *ARF* (one B3 domain, auxin response factor and AUX/IAA domain), *RAV* (one B3 domain and AP2 domain) and *REM* (two B3 domains) have different typical domains [13]. The B3 superfamily plays an important role in plant growth and development as well as stress response [14–15]. ARFs have the function in development of flowers and leaves, vascular tissue differentiation and root initiation [16–17]. In *Arabidopsis thaliana*, *arf1* and *arf2* mutant plant exhibit development delay, including initiation of flowering and rosette leaf senescence [18]. The double mutant *arf7 arf19* influenced the growth of root and leaf expansion [19]. The study of *LAV* is few, it mainly participates in regulation of seed germination, embryo development and stress response [20–23]. *AtABI3* induced lateral root primordia by auxin, the root of loss-of-function *abi3* alleles show insensitivity to auxin [24]. The *LEC2* induces the somatic embryo formation and promotes the accumulation of seed storage protein and oil bodies [25]. As a transcriptional repressor, *HIS* has a function to repress the seed maturation genes [26]. Recently, a study showed *HSI2* repress *AGL15* (relate to seed development) by the PRC2 component MSI1, thereby regulating seed maturation [27]. Many studies showed the *RAV* is related to stress response, for example, overexpression of cotton *RAV1* Gene in *Arabidopsis* increased the sensitivity of salinity and drought stresses [28]. On the other hand, *TEM* (*TEMPRANILLO*) gene (a member of RAV family) downregulate the expression of *FT*, so as to repress flowering [29]. The *REM* is latest studied and little is known about its function. *AtREM34* is the first gene be identified but *VRN1* (*VERNALIZATION1/REM5*) is the first to be functionally characterized, *VRN1* is related to vernalization and promote flowering [6, 30]. Recently, a study silenced *REM34* and *REM35* genes and influenced the female and male gametophyte development [31]. These indicate that *REMs* play roles in flowering.

Pineapple (*Ananas comosus* (Linn.) Merr) is the herbaceous plant of *Bromeliaceae* family. It is one of the four tropical fruits in the world, which is planted in more than 80 countries or regions [32]. There are mainly two ways to induce pineapple flowering: natural flowering induction and artificial flowering induction. Natural flowering induction needs short day-length and cool night temperatures to produce endogenous ethylene, artificial flowering induction use chemicals like ethephon or calcium carbide to release ethylene [33–35]. However, in natural flowering induction, the uniformity of flowering is often inconsistent or pineapples in the planting area cannot be centralized in the market. Artificial flowering induction usually used in commercial production, it can effectively improve flowering regularity and flowering rate, and effectively plan the time to market, so as to achieve the purpose of improving pineapple planting efficiency [36]. There are many genes related to ethylene or flowering which studied in pineapple, but the mechanism of ethylene induces pineapple flowering is unclear.

The articles about B3 superfamily is little by now, just in several plant species, meanwhile, the genome-wide analyses about B3 superfamily in pineapple hasn’t been reported. The members of B3 superfamily play roles on flowering, it’s unclear whether the B3 superfamily genes involve in pineapple ethylene response and flowering. In this study, 57 pineapple *B3* genes were identified. We did comprehensive analysis of gene duplications, chromosome distribution, gene structure and *cis*-acting regulatory elements. Phylogeny were analysed with the *B3* genes of *Arabidopsis* and rice. The expression patterns of *AcB3* genes after ethephon treatment were showed by RNA-Seq data. Expression profiles after ethephon treatment were evaluated by qRT-PCR to analyse the relationship between *B3* and ethephon. This study provided reference for further understanding of the physiological function and mechanism of *B3* genes in ethephon induced pineapple flower formation.

## Materials and methods

### Identification and analysis of *B3* genes in pineapple

To identify *AcB3* genes, The genome of *Ananas comosus* were downloaded from the PGD (Pineapple Genomics database)(http://pineapple.angiosperms.org/pineapple/html/index.html). The HMM (hidden Markov model) matrix of B3 domain (PF02362) was downloaded from Pfam, and then *B3* genes were retrieved by Hmmer 3.0 software. These predicted B3 proteins were further confirmed and analyzed using the CD-search (https://www.ncbi.nlm.nih.gov/Structure/cdd/wrpsb.cgi) and SMART (http://smart.embl-heidelberg.de) web server. All pineapple *B3* genes contained B3 domain.

### Chromosomal locations and synteny analysis

The physical locations of *AcB3* genes were obtained from the PGD. The location images of *AcB3* genes were drawed by MapChart software [37]. Multiple Collinearity Scan toolkit (MCscanX) was used to analyse the tandem and segmental duplications of *B3* genes [38]. The diagrams of synteny analysis were drawed by Tbtools [39]. The value of Ks and Ka were calculated by using DnaSP 5 [40] software.

### Phylogenetic analysis

Multiple sequence alignments and phylogenetic tree of full-length B3 proteins from pineapple, *Arabidopsis* and rice were performed using PhyML 3.0 with maximum likelihood (ML) method, ‘AIC’ criterion and ‘aLRT Chi2-based’ as fast likelihood-based method [41]. The classifications of *AcB3s* were given according to the lists of *AtB3* and *OsB3* genes came from previous study [13]. The sequences of *AtB3* and *OsB3* genes were downloaded from Phytozome12.1.6 (https://phytozome.jgi.doe.gov/pz/portal.html).

### Structural and motif analyses of *AcB3* genes

The exon/intron organizations picture of the *AcB3* genes were drawed by TBtools. The conserved motifs were analyzed using MEME (http://meme-suite.org/tools/meme) [42], searching up to 20 conserved motifs and each motif was set from 6 to 50 residues. Protein domains were identified using the NCBI CD-search and SMART.

### Analysis of *cis*-acting elements and gene ontology

The 2kb promoter region upstream of the start codon of each gene was downloaded from PGD to analyse the possible *cis*-acting elements of *B3* genes by PlantCARE (http://bioinformatics.psb.ugent.be/webtools/plantcare/html/) online server. WEGO (http://wego.genomics.org.cn/) was used to analyse Gene ontology of B3 proteins.

### Plant materials and treatments

The pineapple hybrid variety was "Tainong 16" is cultivated in the pineapple planting base of Chengmai County, Hainan Province, China. The temperature, humidity and other environmental conditions in the greenhouse which controlled in the appropriate range for pineapple flowering. The pineapple which growed 25 mature leaves(more than 30 cm in length) were treated by 300 fold solution of 40% Ethephon (1600 mg/L), and were sampled at 1h, 3h, 6h, 1, 6, 10, 15, 22 and 31 days after treatment, including leaves, stem apex. The sample treated with water of the same volume as the contrast, all of them were frozen in liquid nitrogen and stored in refrigerator at −80^℃^C until used.

### RNA isolation and *AcB3* genes expression analysis

Total RNA was extracted from samples using Total RNA was extracted from samples using a universal plants RNA extraction kit for Hua Yueyang, according to the manufacturer’s guidelines. 1 μg RNA were used to synthesize the first-strand cDNA by MonScript^™^ RTIII All-in-One Mix with dsDNase. Primers were designed for real-time quantitative PCR (qRT-PCR) using Primer 5.0 software, and the primer sequences were shown in detail in S7 Table. The cDNA was used as the template, *AcEF1* as the internal reference gene, and each sample was repeated three times for real-time fluorescence quantitative PCR analysis. The reaction was carried out on Roche Lightcyler^®^ 480 instrument using SYBR Green Master Mix (Roche). The reaction system of real-time quantitative PCR was 10 μL, including 1 μL of cDNA, 5 μL of SYBR Green I master, 0.5 μL of 10 μmol/L of forward and reverse primers, and 3 μL of ddH_2_O. The data were analyzed by 2^−ΔΔ CT^ method [43], and by Excel and Graphpad prism 7.0 softwares. The transcripts of the similar sequences were found by local blast, and the fragments per kilobase of exon model per million mapped reads (FPKM values) were calculated, finally the heatmap was drawn by pheatmap software base on the log_2_ (FPKM+1) values. The transcriptome data were uploaded to NCBI (PRJNA639649).

## Results

### Identification and characterization

To comprehensively identify the candidate B3 proteins in pineapple, 62 *AcB3* genes were found by HMMER 3.0. Removing repetitive (Aco026957 and Aco017311) and short sequences (Aco017668 and Aco017067), the CD-search and SMART analyses ware used to verify the presence of the B3 domains later, one (Aco015073) didn’t contain B3 domain. Finally, 57 pineapple B3 proteins were identified.

Gene characteristics, including the protein properties, protein secondary structure and and the subcellular localization were analyzed (S1 Table). Among the 57 B3 proteins, the longest protein is 997 aa and the shortest is 137 aa with molecular weights (MWs) from 15.71KDa to 111.32KDa. The isoelectronic point (PI) values ranged from 4.61 to 9.9, all of them are hydrophobic proteins. The prediction of subcellular localization shows they are mainly located in nuclear (42), chloroplast (8), cytoplasmic (5), extracellular (1), vacuole (1).

### Genomic location and gene duplication analysis

Chromosomal location analyses (Fig 1) showed that 54 pineapple B3 genes were distributed on 21 out of the 25 Linkage Groups (LG), the rest (Aco030005, Aco030007 and Aco030008) were located on scaffold_1004. The distribution was uneven, LG14 contained the largest number of 8 B3 genes followed by 7 on LG01 while only one B3 gene was found on LG5, 6, 8, 9, 12, 13, 15, 18. Seven genes were highly concentrated on 9.94Mb to 10.24Mb of LG14.

**Fig 1.**
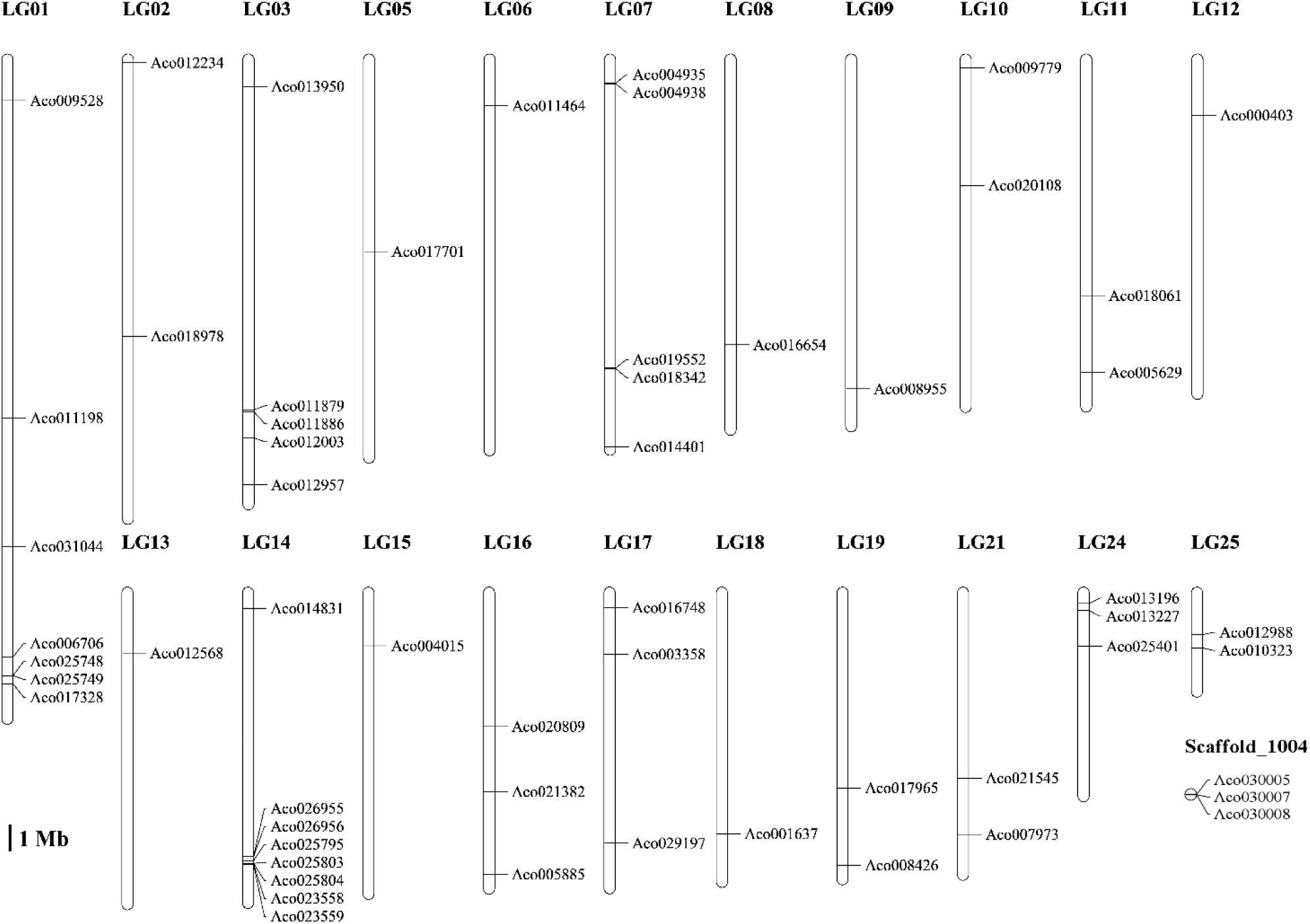
Distribution of *AcB3* genes on linkage groups. All 57 *B3* genes are mapped onto 21 linkage groups and one scaffold. “LG number” indicated the chromosome number. The scale is in megabases (Mb).

To investigate the potential duplication events of *B3*, both BLASTP and MCScanX method were used to identify the collinearity of the *B3* gene family in pineapple. 8 segmental duplication events were identified (Fig 2), and the synteny blocks of *B3* were on 12 LGs (LG01, 03, 05, 11, 12, 14, 15, 16, 17, 19, 21 and 24). 5 tandem duplication events were also identified in LG14 and scaffold_1044 (S2 Table). Calculating the nonsynonymous (Ka) and synonymous (Ks) substitution rates is useful for the study of evolutionary, Ka/Ks=1, indicates neutral mutation; Ka/Ks>1, indicates negative (purifying) selection; Ka/Ks<1, indicates positive (diversifying) selection.In these 13 duplication *B3* gene pairs, 2 of them were “Ka/Ks>1”, implying that those had evolved under the effect of positive selection; 10 of them were “Ka/Ks<1”, implying that those had evolved under the effect of purifying selection; the only one pair had Ks value equal to 0 and the Ka value is very small, implied they may be the redundant gene.

**Fig 2.**
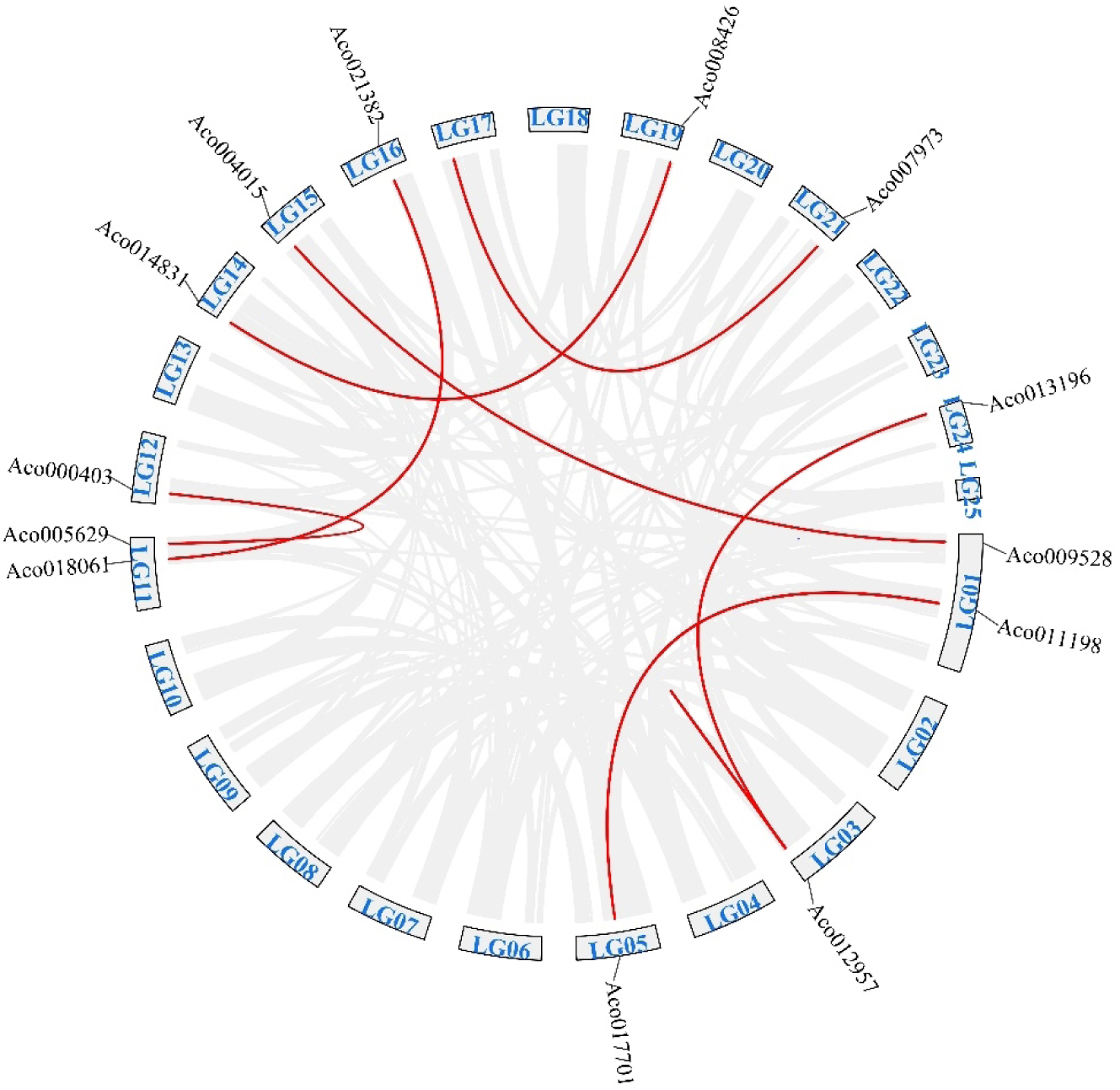
Collinearity analysis of *B3* genes in pineapple. Gray lines suggest all segmental duplications and the red lines suggest duplicated *B3* gene pairs. The chromosome number and gene ID are showed.

To further study the potential evolutionary mechanisms of the pineapple *B3* genes superfamily, the collinear relationship between pineapple and other two species *Arabidopsis* (dicotyledon) and rice (monocotyledon) was analyzed (Fig 3). We identified 7 collinear *B3* gene pairs between pineapple and *Arabidopsis*, 39 collinear *B3* gene pairs between pineapple and rice. The details were showed in S3 Table. The number of orthologous events of *AcB3s*-*OsB3s* was far more than that of *AcB3s*-*AtB3s*, which indicated the genetic relationship between pineapple and rice is closer than that *Arabidopsis*.

**Fig 3.**
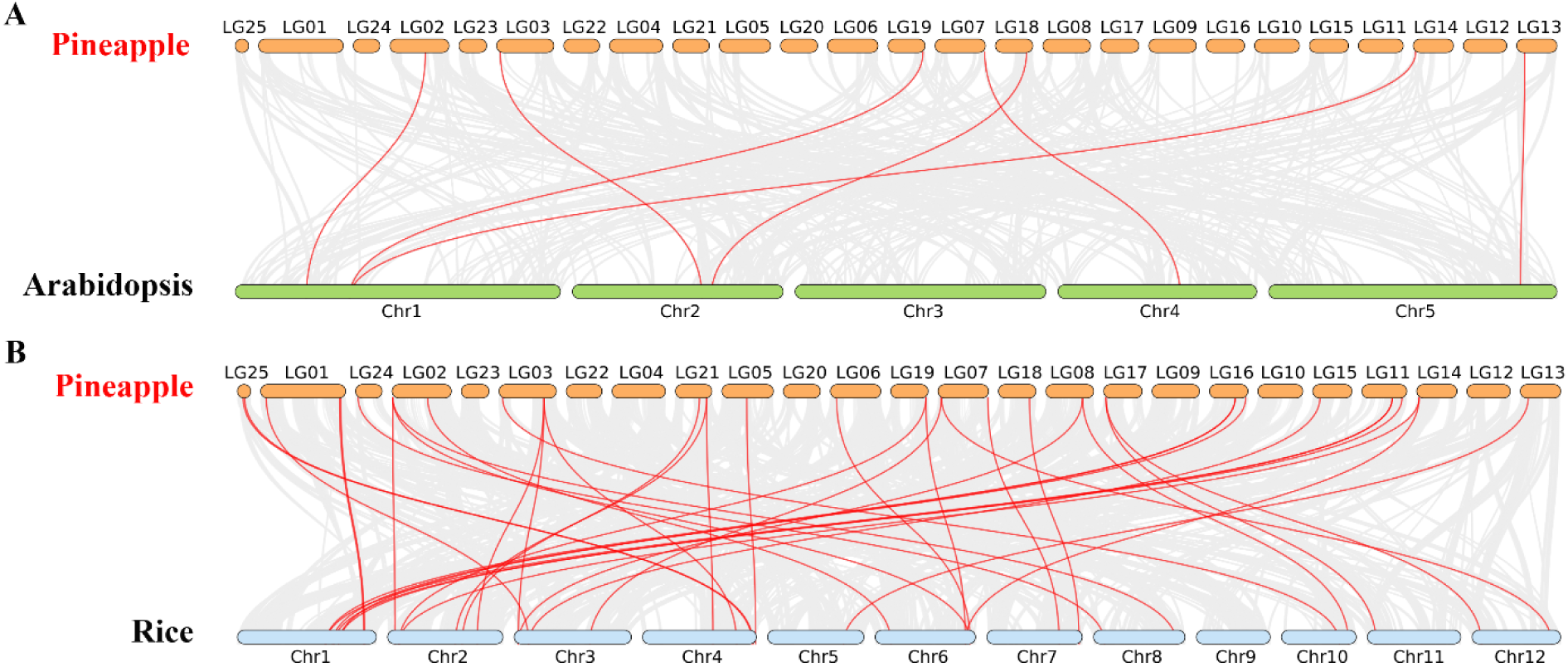
Collinearity analysis of *B3* genes between pineapple, *Arabidopsis* and rice. Gray lines indicated the collinear blocks between pineapple and two other plants, the red lines meant the syntenic *B3* gene pairs.

### Phylogenetic analysis among the pineapple, *arabidopsis* and rice B3 proteins

To study the evolutionary relationship of B3s among pineapple, *Arabidopsis* and rice, we constructed a phylogenetic tree using MEGA 6.0, an unrooted phylogenetic tree was created. In this study, 87 B3 proteins from *Arabidopsis* [13], 86 from rice [13], 57 from pineapple were utilized for the phylogenetic analysis.

As shown in Fig 4, these proteins can be divided into five distinct groups. 27 *AcB3* genes were belonged to *REM*, making it the largest subclass, followed by 19 *AcB3* genes belonged to *ARF*, *RVA*, *HSI* and *LAV* subfamily had 6, 3 and 2 *AcB3* genes, respectively.

**Fig 4.**
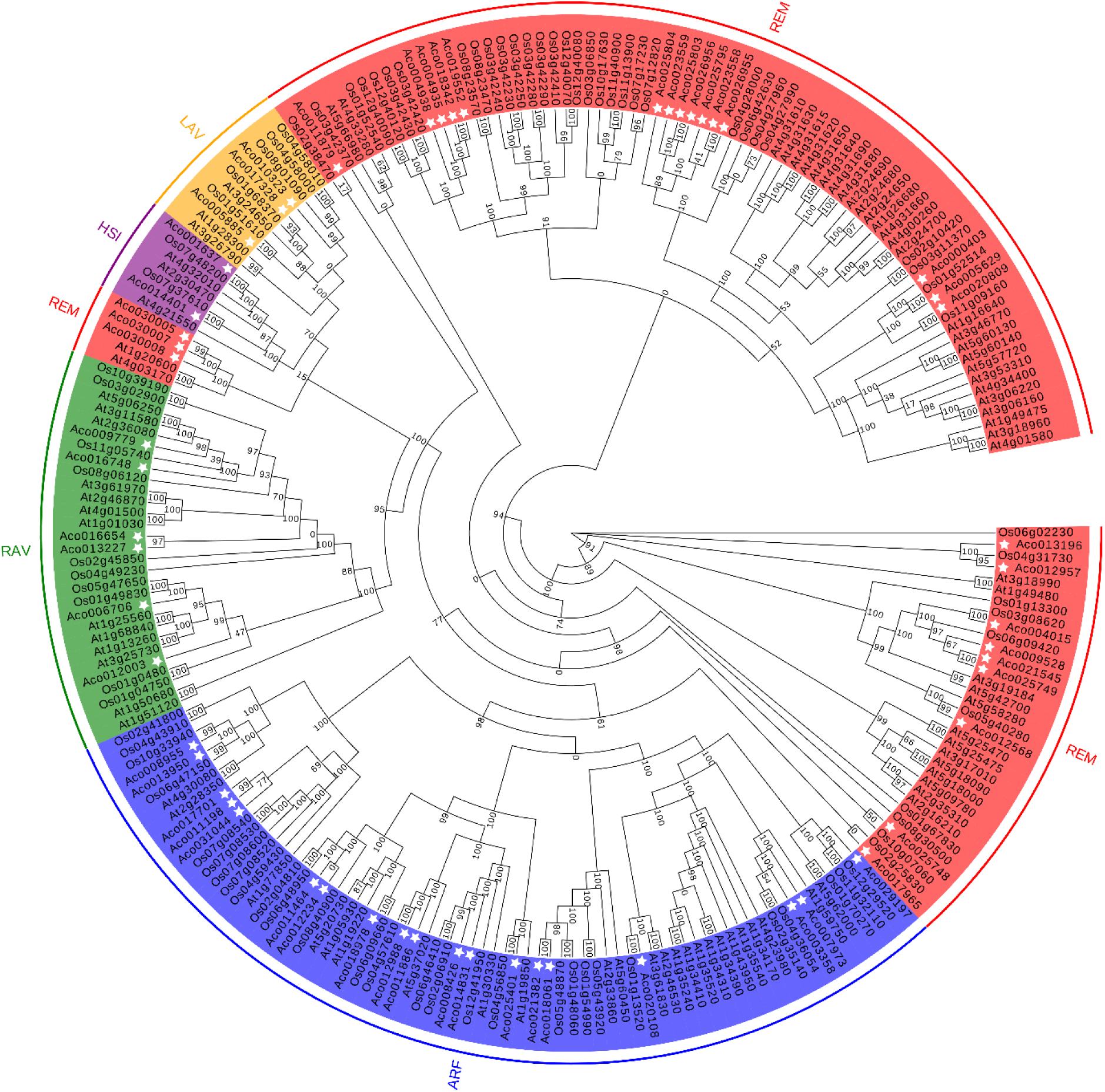
The phylogenetic tree of B3 proteins from pineapple (Ac), *Arabidopsis* (At) and rice (Os). The different color area indicated different groups, the white asterisk indicated *B3* genes from pineapple.

### Structural and motif analysis

To investigate the structural diversity of *AcB3* genes, the structures of *AcB3* genes were analysed. 57 *B3* genes were divided into 5 groups according to Fig 4. Different subfamily had different number of exons, ranging from 2– 15 and 1–8 in ARF and REM families, respectively. Most ARF subfamily genes had more than 11 exons and more than half REM subfamily genes had 3-5 exons. The LAV subfamily genes have 6-8 exons, the HSI subfamily genes have 12 or 13 exons, and 1 or 2 exons in RAV subfamily genes (Fig 5B).

**Fig 5.**
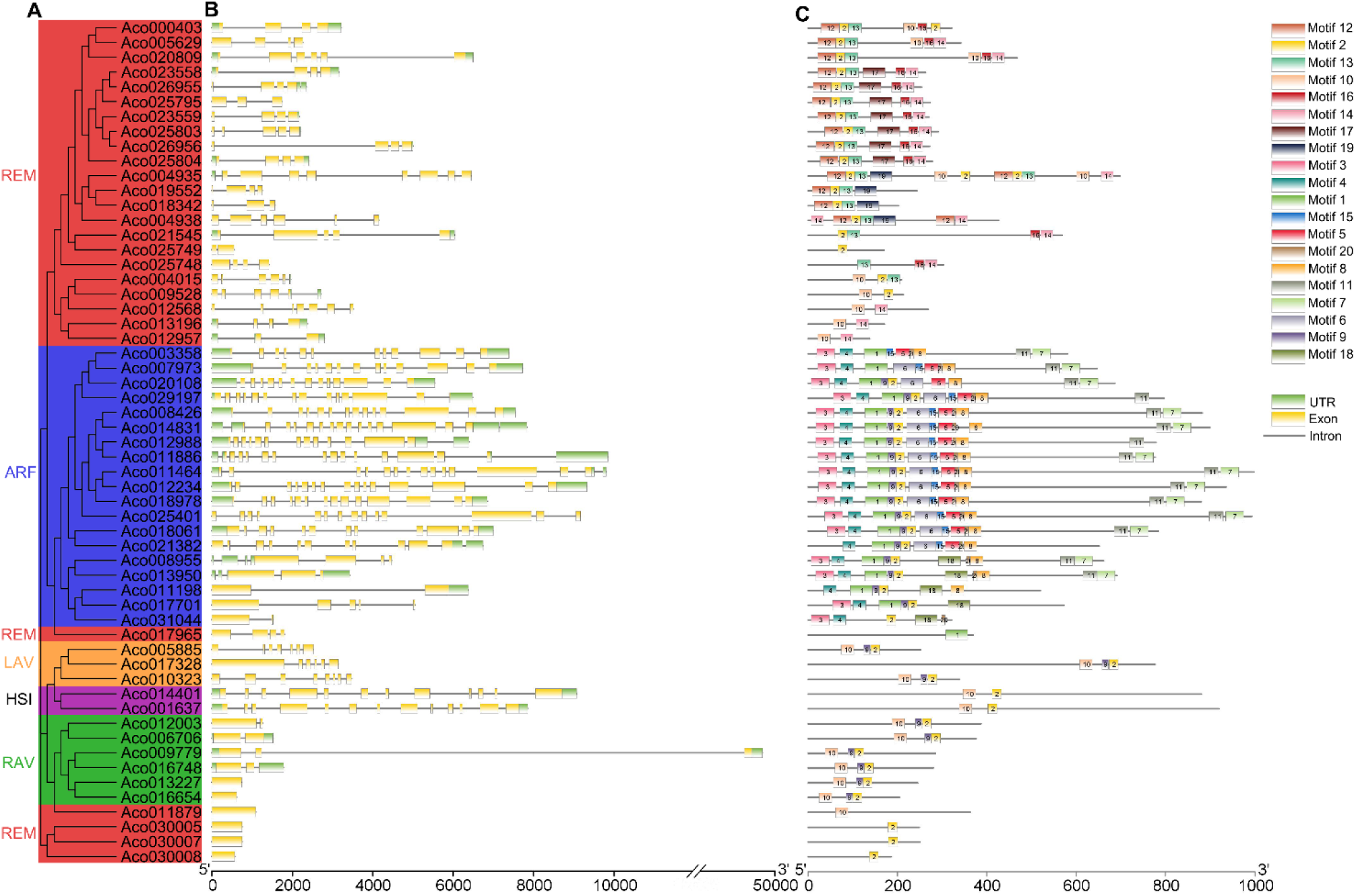
The exon-intron structure and motif organization of *AcB3* genes. (A) Phylogenetic relationships of *AcB3* genes. (B) The intron and exon structure of *B3* genes. The green and yellow boxs represent exon and UTR, respectively, the line which go through the boxs represent intron. (C) The motif identification analysis of AcB3 proteins using MEME.The detailed information of 20 motifs are listed in S4 Table.

To investigate the protein sequence features of *AcB3s*, we identified 20 different motifs (Fig 5C). Most motifs are 28-50 amino acids in length, and the details of 20 motifs were presented in S4 Table. As the result showed all *AcB3* genes had motifs which annotated as B3 domain. The same group had the similar motifs, while motifs were diverse among different groups. For example, both *HSI* genes had motif2 and 10, while all *ARF* genes had motif3 and 4. Meanwhile, the motif12, 13, 14 16, 17 and motif19 were only existed in *REM* (Fig 5C). The special motifs in groups may imply diverse functions of AcB3 superfamily.

### *Cis*-acting elements and gene ontology analysis of *B3* genes

To investigate the potential regulatory role of B3 genes, we analysed promoters which were 2000bp in length before ATG, The *cis*-elements related to plant hormones were showed in Fig 6. There were 42, 26, 40, 28, 35, 15 *AcB3* genes promoters had *cis*-elements related to abscisic acid, auxin, ethylene, gibberellin, methyl jasmonate and salicylic acid, respectively. Aco009528 and Aco000403 didn’t contain these 6 plant hormone related elements. Other *cis*-elements such as light responsiveness element, defense and stress responsiveness element and so on were also found, more detials were showed in S5 Table.

**Fig 6.**
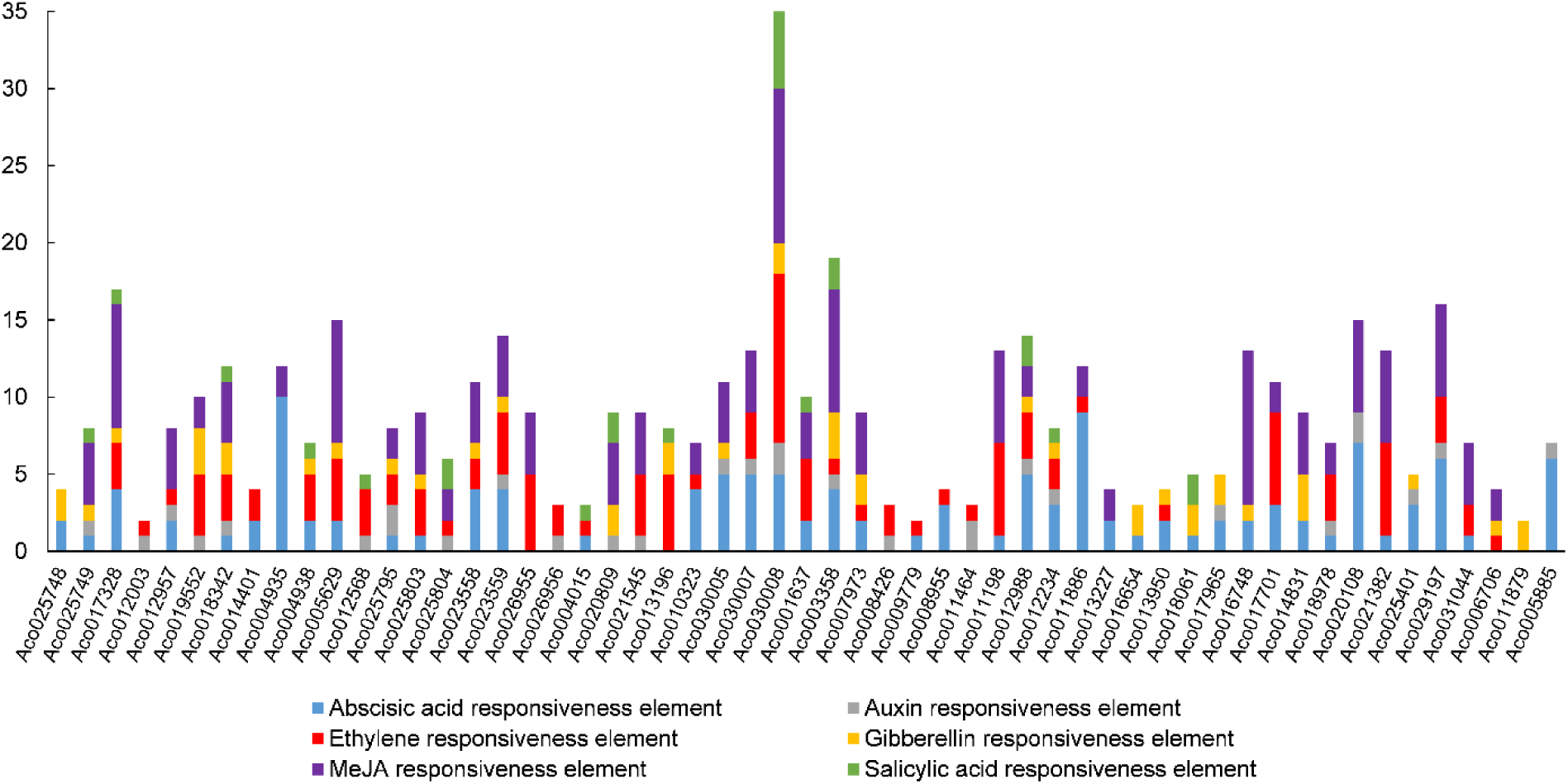
B3 promoters *cis*-elements prediction of hormone related. The promoter is 2.0 kb in length.

We also executed a gene ontology (GO) enrichment analysis of the *AcB3* genes(Fig 7). The prediction of cellular component suggested *AcB3s* participated in cell, organelle, protein-containing complex. The prediction of biological process suggested most *AcB3s* participated in DNA binding then transcription regulator or catalytic activity. Moreover, the prediction of molecular function suggested *AcB3s* mainly involved in metabolic process, cellular process, regulation of biological process and biological regulation. The terms on level 3 were also analysed (S1 Fig; S6 Table), significantly, there was two genes (Aco13950 and Aco006706) involved in the function related to flowering.

**Fig 7.**
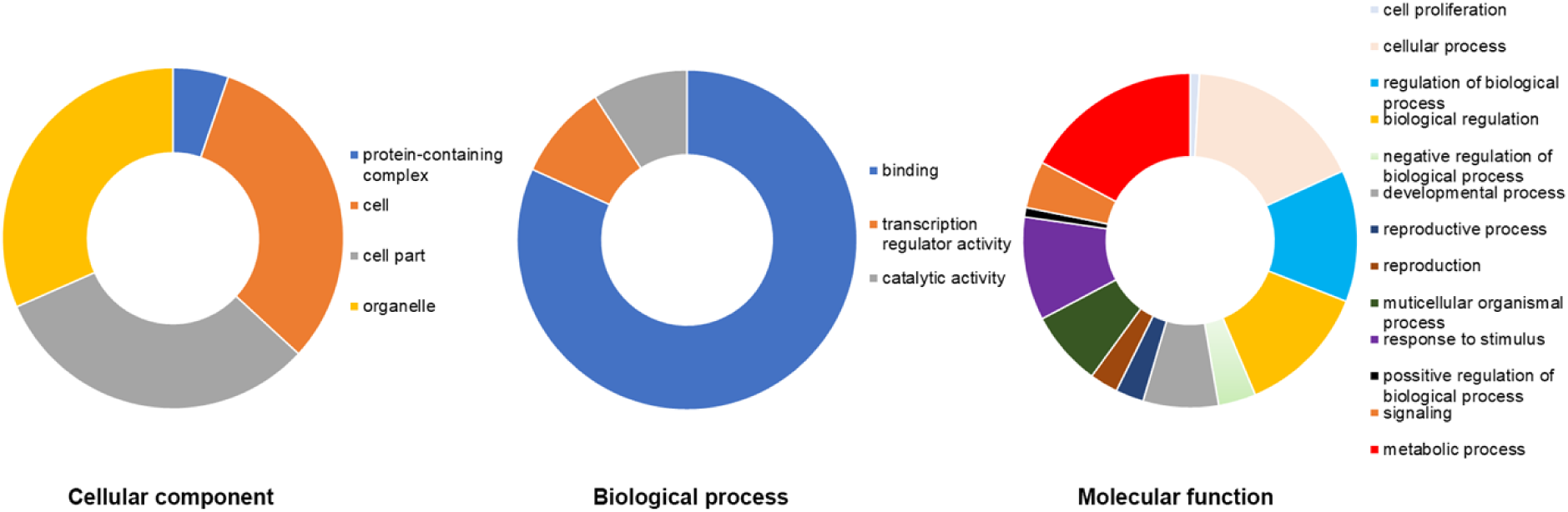
Gene ontology analysis of AcB3 proteins. Three categories (cellular component, biological process and molecular function) and terms on level 2 were showed by different color.

### Expression profiles of *AcB3* genes in stem apex after ethephon treatment

To investigate if *AcB3* genes were regulated by ethephon, the FPKM value of 47 *AcB3* genes after ethephon treatment were collected from RNA-Seq data and presented in a heatmap. The transcripts of 10 *AcB3* genes were not detected in RNA-Seq data, indicated these may expressed in other growth period or tissues. As showed in Fig 8, more than half of *AcB3* genes were highly expressed, 16 *AcB3* genes were up-regulated compared with CK and 6 *AcB3* genes were down-regulated compared with CK. Aco018978, Aco017965, Aco018061, Aco012988, Aco011886, Aco001637 and Aco014831 showed higher level of expression after ethephon treatment. These findings indicated that *AcB3* genes expressed in response to ethephon.

**Fig 8.**
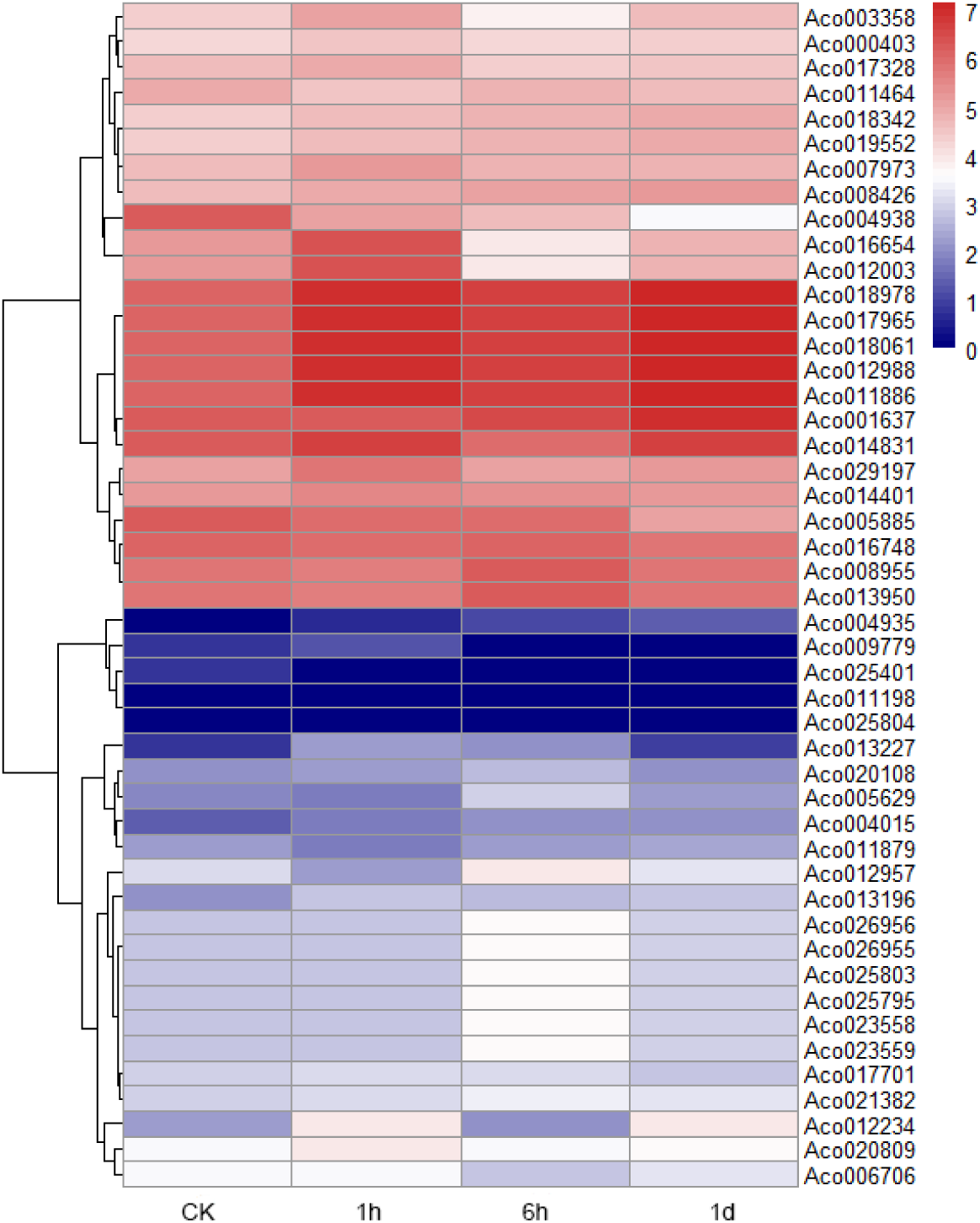
Heatmap of *AcB3* genes after ethephon treatment in stem apex. The color scale is based on the log2 (FPKM + 1) values

### Expression analysis of *AcB3* genes after ethephon treatment

We found 31d after ethephon treatment in pineapple, the flower bud differentiation was in progress in previous study [44]. In order to identify *AcB3* genes resopnsing to ethylene and the functioning on flowering, 20 genes from different subfamilies were randomly selected to analyze their expression patterns by qRT-PCR. In general, the expression levels of all the selected genes were significantly changed in response to ethylene. In leaves, most genes were up-regulated, while Aco010323, Aco001637, Aco018978 and Aco025401 were significantly down-regulated through all the time points. The expression levels of some genes like Aco004938, Aco026955, Aco020809 and Aco021545 were up-regulated before 1d, and the expression levels of 8 genes were significantly up-regulated in 31d (Fig 9).In stem apex, the of all 20 genes were up-regulated to different degrees. During 0 to 6d after ethylene treatment, all 20 genes were up-regulate, 8 genes reached peak value at 3h, while 11 genes at 6h and 1 gene at 1d (Fig 10). what’s more, the expression levels of all 20 genes except Aco012957, Aco020809 and Aco013196 were significantly up-regulated in 31d. In particularly, the expression levels of Aco012003 and Aco004938 were at high level all the time and Aco019552 had a higher level of expression among 20 genes, which indicated these *AcB3* genes may involved in flower bud differentiation.

**Fig 9.**
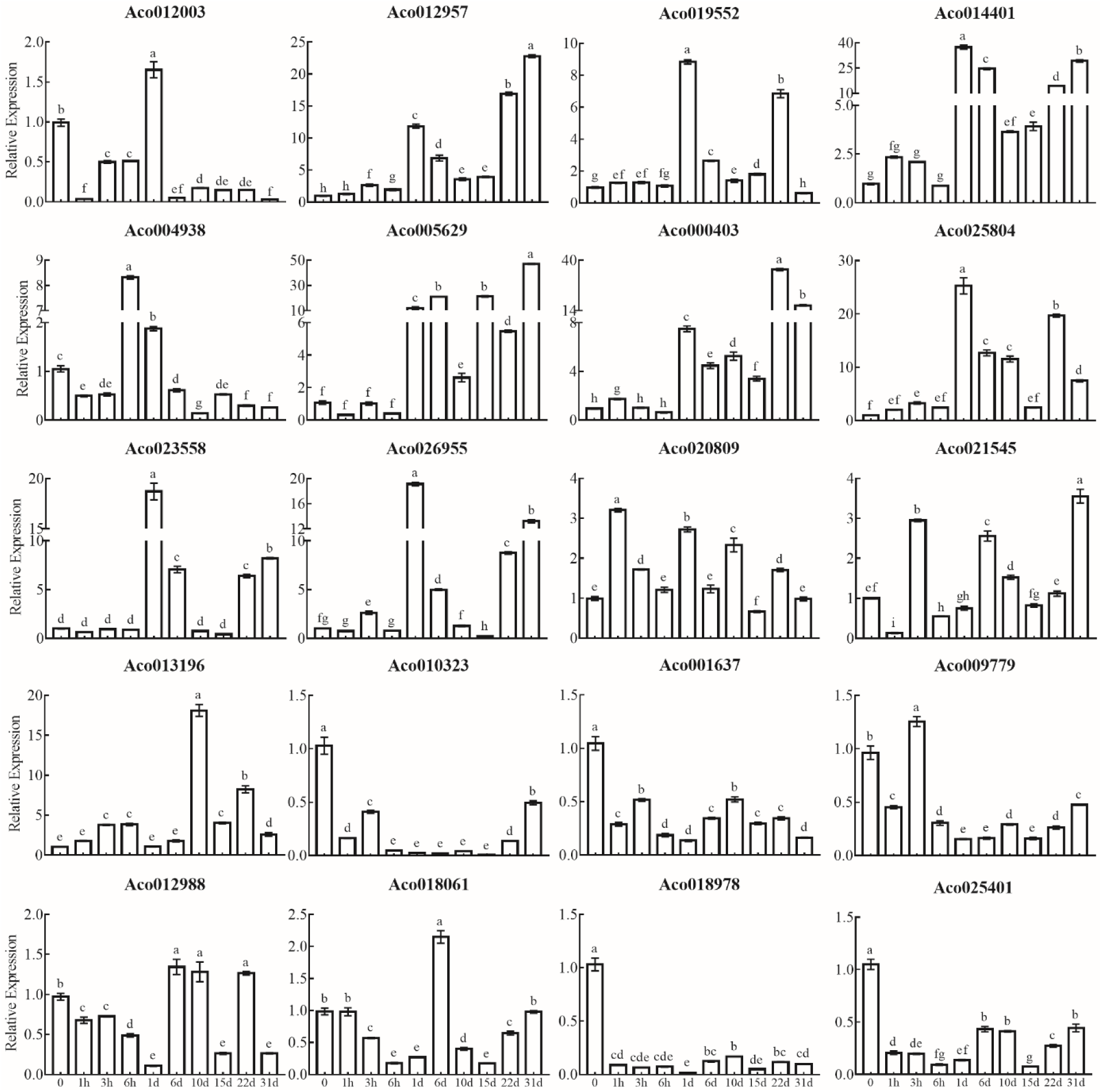
Expression analysis of *AcB3* genes after ethephon treatment during floral induction period in leaves. The ‘0’ was regarded as a standard, and *AcEF1* gene was used as internal control. Error bars indicate standard error (SE) based on three replicates. Different letters indicated significant difference, LSD test (*P*≤0.05).

**Fig 10.**
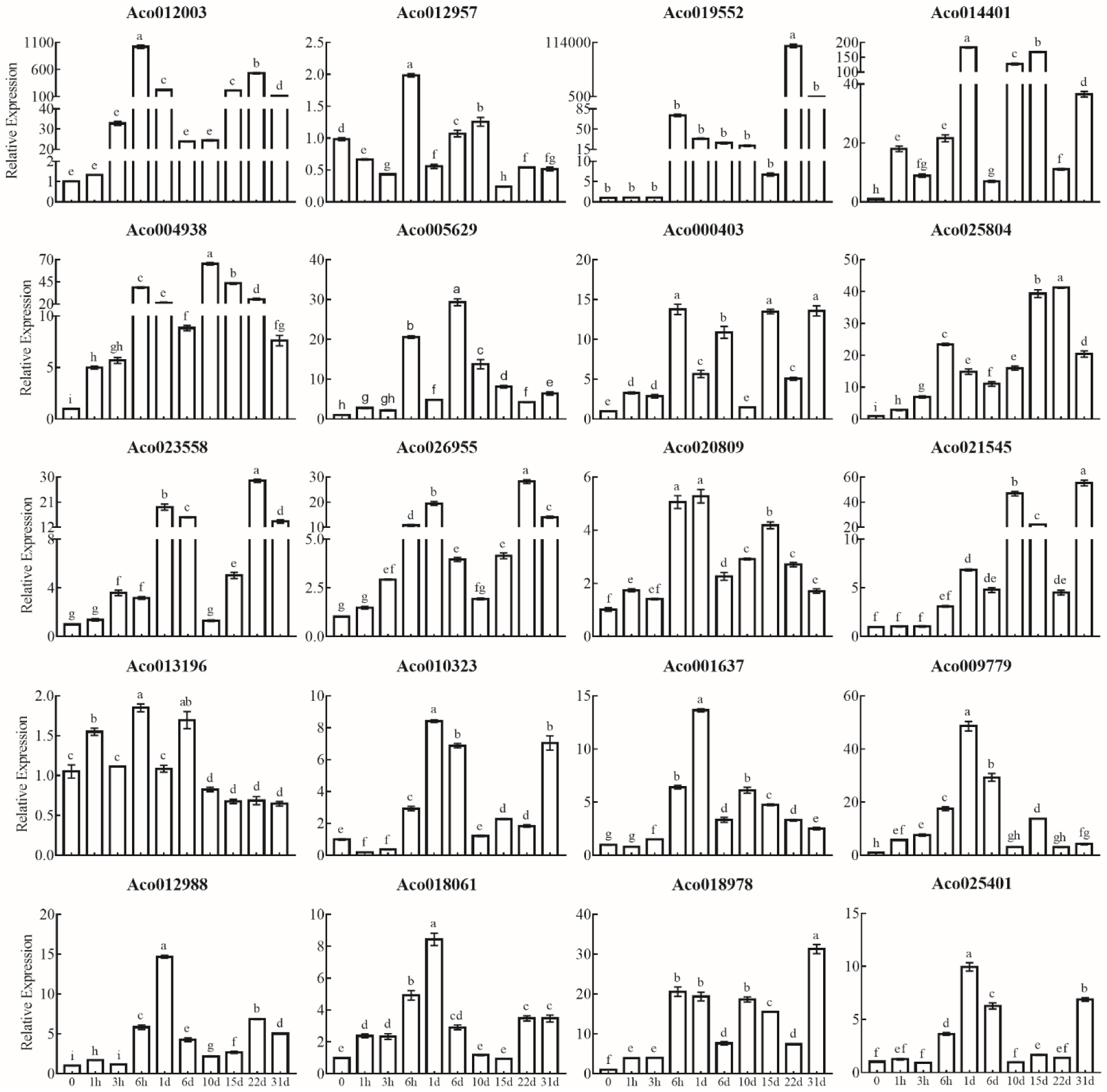
Expression analysis of *AcB3* genes after ethephon treatment during floral induction period in stem apex. The ‘0’ was regarded as a standard, and *AcEF1* gene was used as internal control. Error bars indicate standard error (SE) based on three replicates. Different letters indicated significant difference, LSD test (*P*≤0.05).

## Discussion

### Characterization and structure of the pineapple *B3* genes superfamily

*B3s* had various functions include regulating on root, seed and flower development, responses of organisms to abiotic stresses, involving in hormone signaling pathways [7, 12]. Genome-wide characterization and expression analyses about B3 superfamily had been investigated in some plants [8, 15, 45–46]. In this study, a comprehensive genome-wide analysis of the AcB3 superfamily in pineapple was performed for the first time and 57 *AcB3* genes were identified. The number of *AcB3* genes were less than the other species (*Arabidopsis* and rice), but had same five subfamilies, which indicates that parallel evolutionary events of *B3* genes existed in dicots and monocots.

Different gene structures have different function and the introns have impacts on genes expression and evolution [47]. In this study, most *ARF*, *LAV*, *REM* and *RAV* genes had 3, 4, 1 or 2 and 0 introns in B3 domain, respectively. This number in HSI was 2. These indicated similar B3 structure existed in same subfamily during the evolution process. Although 4 motifs were annotated as B3 domain (S4 Table), all 57 *AcB3* genes had B3 domain, this supported the characterization of *B3* genes. Similar protein architecture may have similar function, in this study, most genes had similar classification and number of motifs in the same subfamily, such as *REM* and *ARF*. Together, these results were similar with those of the phylogenetic analysis, which supported the reliability of our subfamily classification.

### The evolutionary analysis of the *AcB3* genes superfamily

In this study, 57 *AcB3* genes were classified into five subfamilies by phylogenetic analysis, the same subfamily has similar domains (S2 Fig). Like other species [9], the REM subfamily had the most genes followed by ARF subfamily. Previous study showed, there were 20 *ARF* genes in pineapple [48]. But in this study, we identified 19, the gene (Aco015073) didn’t contain B3 domain. Within the ARF subfamily in this study, 14 of them contained B3, Aux_resp and Aux_IAA domains, the other 5 have B3 and Aux_resp domain. This indicated during the evolutionary process the B3 and Aux_resp domains were more conservative than the Aux_IAA domain. The RAV subfamily usually had one AP2 and one B3 domain, but not all RAV protein contained AP2 domain [13, 49]. In this study, Aco009779, Aco016748, Aco013227 and Aco016654 don’t contain AP2 domain grouping with RAV genes from rice and *Arabidopsis* belonged to RAV subfamily. The REM subfamily had B3 domains from 1 to 4, multiple B3 domain may be caused by domain duplication events.

Tandem duplication, segmental duplication, and transposition were the mainly gene duplication events, these events promoted the expansion of gene families, and segmental duplication had more proportions than tandem duplication [50–51]. In this study, we identified 8 segmental duplication events and 5 tandem duplication events, most had the Ka/Ks value less than 1, indicated positive selection happened in these genes. Most genes were from REM subfamily, one pair belonged to ARF subfamily (Aco018061 and Aco021382) which Ka/Ks value was more than 1, they may have different function. Synteny analysis were also carried out with two other species, *Arabidopsis* and rice, which belonged to dicot and monocot plants, respectively. Syntenic gene pairs between pineapple and *Arabidopsis*, rice were 7 and 39, respectively. This indicated pineapple and rice had closer evolutionary distance than pineapple and *Arabidopsis*, which was consistent with previous publications [52–53]. Moreover, genes in the same Syntenic gene pairs between pineapple and rice may had similar structure and function.

### *Cis*-element analysis in the promoters of *AcB3s*

*Cis*-element was short in length with a core of only seveal base-pairs, but could regulate gene expression and related to the functions of downstream genes [54–55]. The results illustrate that 6 phytohormones-related *cis*-element were existed in *AcB3* genes superfamily, include abscisic acid, auxin, ethylene, gibberellin, MeJA and salicylic acid responsiveness elements. Most *AcARFs* (17 of 19), both *AcHSIs* and all *AcLAVs* had abscisic acid responsiveness elements, and most *AcREMs* had abscisic acid, ethylene and MeJA responsiveness elements, which indicated they were participated in response to abscisic acid and abiotic stress. Pineapple is a special plant, which ethylene could promote flowering [56]. There were 40 *AcB3s* had ethylene responsiveness elements, 26 *AcB3s* had auxin responsiveness elements which related to development, but only 19 *AcB3s* had both elements, these genes may involve in the process of pineapple flowering induced by ethylene. The other elements were showed (S5 Table), include anaerobic induction, circadian control, defense and stress, drought inducibility, light and low temperature responsiveness element. Most *AcB3s* (52 of 57) had light responsiveness element followed by anaerobic induction element (44 of 57 *AcB3s*). These results suggest that *AcB3s* had various functions and researchs shoud be carried out for more evidences.

### The potential roles of differentially expressed *AcB3s*

To investigated gene functions, the gene expression patterns could provide important reference. Previous study showed, *FT* gene which overexpression caused early flower, showed high level of expression in stem apex after ethephon treatment and the time when flower bud differentiation was in progress. In this study, we found all 20 *AcB3* genes had responsiveness to ethylene in stem apex within 1d, only 17 *AcB3* genes in stem apex showed significant up-regulation in 31d. This indicated these may play roles in flower development.

The studies of REM subfamily genes were few, these genes involved in flower development and most of them preferentially expressed in reproductive meristem and were the targets of the key floral transcription factors like *AP1*, *AP3*, *PI*, *LEAFY* and *SVP* [6, 57–58]. In *Arabidopsis*, *AtREM22* was related to stamen development and *AtREM5* (*VRN1*) was involved in floweing time control [59–60]. In this study, compared the expression in leaves and stem apex, the relative expression of most *AcREMs* in stem apex were higher than leaves, like *REMs* in other species. High level expression of *AcB3s* in stem apex and especially in 31d indicated these genes may have functions in flowering. *AtREM17* (At4g34400) expressed preferentially in flower and was a target of *LEAFY* [6, 8], Aco000403, Aco005629 and Aco020809 have a close genetic distance with At4g34400, they may have similar function in flower development.

It was reported that HSI mainly regulated seed maturation [27], but there was exception. *AtVAL1* (At2g30470) also named as *AtHSI2* which was the target of *FLC* (*Flowering Locus C*) and may play roles in flowering process [61]. Aco001637 changed largely in 6h after ethephon treatment and expressed high level in 31d, this gene was classified together with At2g30470 by phylogenetic analysis, indicated similar function in pineapple.

Auxin has important roles on plants growth and development, *ARF* genes could response to auxin and were best studied [62]. ARFs involved in flowering process, for example, *AtARF3* regulate floral meristem determinacy by *AG* and *AP2* [63]. In tomato, most *ARFs* had higher expression level in flower buds and flower compared in fruit [64]. In this study, *ARFs* showed low relative expression in leaves but high in stem apex, all of them up-regulated significantly in 31d, indicated they may involve in flowering process. Moreover, all of them up-regulated significantly in 6h after ethephon treatment, this suggested they may response to ethylene. *Arabidopsis ARF6* and *ARF8* (At5g37020) coordinate the transition from immature to mature fertile flowers [65], Aco012988 classified together with At5g37020 may has similar function. Phosphorus was necessary for plant growth and development, the concentration of phosphorus had influence on flowering [66]. To adapt low phosphorus environment, ethylene can adjust the relative elongation of roots and change the signal transduction pathway of phosphoric acid [67]. As a gene of ARF family, *OsARF16* (Os06g09660) had diverse function involved in phosphate transport [68]. Intriguing, Aco018978 was classified together with Os06g09660, Aco018978 up-regulated significantly after ehephon treatment and in 31d, suggested that ethylene may increase the length of root and high expression level of Aco018978 may increase phosphate transport to the flower formation in the future.

## Conclusions

In this study, 57 *AcB3* genes were identified in the pineapple genome for the first time. They are unevenly distributed on the chromosomes, and segmental duplications contributed to the expansion of the AcB3 family. It was also found that *B3* genes from pineapple and rice have closer genetic distance. Phylogeny analysis comparing with *Arabidopsis* and rice classifies these genes into five major classes (*REM*, *ARF*, *RAV*, *HSI* and *LAV*). *Cis*-element analysis discovered several phytohormone and stresses responsive elements in the promoter region of the *AcB3s*. Moreover, most selected *AcB3* genes express in response to ethylene and show high level expression in flower bud differentiation period, which disclosed that these have roles in flowering and responsing to ethylene signal. Our research provides new insight into the evolution and divergence of *AcB3* genes and helps in identifying candidate genes for functional characterization in flowering. Furthermore, these results will also serve as a foundation and reference for future research regarding the molecular mechanisms in ethylene induced pineapple flowering.

## Author Contributions

Conceptualization: Cheng Cheng Ruan, Zhe Chen.

Methodology: Cheng Cheng Ruan, Zhe Chen.

Resources: Xianghe Wang, Li Jun Guo.

Software Formal analysis: Hong Yan Fan, Fu Chu Hu.

Visualization: Zhi Wen Luo.

Writing - Original Draft: Cheng Cheng Ruan.

Writing - Review & Editing: Zhi Li Zhang.

All authors approved the final manuscript.

